# Deeplearning based MHC epitope prediction for cancer neoantigen discovery

**DOI:** 10.1101/2021.11.10.468160

**Authors:** Carolyn Xie, Yu Shi, Chi Zhang

## Abstract

Neoantigens are important for cancer immunotherapies or cancer vaccine development, but identification of neoantigens is challenging. The high binding affinity between the mutated peptide and MHC (major histocompatibility complex) molecules of the patients is a necessary factor for a somatic mutation on the tumor genome to form a neoantigen. MHC epitope prediction tools can be used for the identification of neoantigens. This research investigates MHC epitope prediction by utilizing Tri-peptide similarity as features for the XGBoost classifier. This model was tested on experimentally validated cancer neoantigen peptides.

## Introduction

Neoantigens, mutated peptides in cancer cells, are derived from somatic mutations absent from normal tissues, and can trigger anti-tumor T-cell responses. A key trait of neoantigens to consider is the fact that they contain a higher binding affinity with MHC (major histocompatibility complex) molecules and therefore are not affected by central immunological tolerance (Peng et al., 2019). Neoantigen-based cancer vaccines in early clinical trial development have been shown to provide host immunity against tumor cells, a significant discovery as they are capable of inducing anti-tumor immune responses through T-cell mediated cytotoxicity (Blass and Ott, 2021; Jiang et al., 2019; Peng et al., 2019). Although neoantigens are important for cancer immunotherapies, identification of neoantigens is challenging. The high binding affinity between the mutated peptide and MHC (major histocompatibility complex) molecules of the patients is a necessary factor for a somatic mutation on the tumor genome to form a neoantigen (Peng et al., 2019).

Many MHC epitope prediction tools, such as NetMHCpan (Jurtz et al., 2017), NetMH-CIIpan (Jensen et al., 2018), MHCflurry (O’Donnell et al., 2018), ConvMHC (Han and Kim, 2017), PLAtEAU (Alvaro-Benito et al., 2018), and NetCTLpan (Stranzl et al., 2010), were developed. However, existing neoepitope prediction methods still have a low rate of validation (Bjerregaard et al., 2017) because the training datasets, usually from IEDB (Vita et al., 2015), were not obtained by standardized experimental methodologies in cancer models (Martins et al., 2019).

In this work, we filtered previously published datasets to keep 9 amino-acid long peptides that have a high binding affinity with the human MCH-I class (i.e. HLA-A, HLA-B, and HLA-C). In developing the MHC binding prediction, the deeplearning model, XGBoost (Chen and Guestrin, 2016), was used. XGBoost is a specialized machine learning model, which provides a gradient boosting framework. XGBoost is popular in classifying and predicting the tabular dataset with superior performance. Tri-peptide similarities, which were used for B-cell epitope prediction (Yao et al., 2012) but not for MCH epitope prediction, were employed as features for the classifier. This model was tested on independent cancer neoantigen datasets, which are experimentally validated cancer neoantigen peptides. If comparing the cancer mutant peptides and corresponding wildtype peptides, our model has success rates from 52.2% to 72.2%, and the corresponding area under receiver operating characteristic curve (ROC) is 0.685.

## Results

### Prediction performance on the training dataset

With the five-fold cross validation, we obtained the average AUC of 0.728176 with a standard deviation of 0.013953. Then, the entire training dataset was used to train the model, and this model was used for independent test datasets.

### Classification performance of neoantigens

Our model was trained with all data in the training dataset, and we applied this classifier model to four cancer neoantigen datasets. We calculated the prediction scores for each cancer mutant peptide and its corresponding wildtype peptide. The difference between the cancer mutant peptide and the corresponding wildtype peptide is only one amino acid mutation in all nine amino acids. If the prediction score of the cancer mutant peptide is higher than the prediction score of its corresponding wildtype peptide, we consider the prediction model is successful. Table 1 shows the success rate for these four datasets.

**Table 1.** Independent tests on known cancer neoantigens.

|  | HC-neoantigenes | MC-neoantigenes | NEPdb | IEDB |
| --- | --- | --- | --- | --- |
| Success rate | 72.2% | 63.2% | 63.3% | 52.2% |

We also merge all cancer mutation peptides from four datasets as a positive dataset, and randomly collected a similar number of negative peptides from NEPdb (Xia et al., 2021) as negative sets. We applied our classifier model to this dataset as well, and AUC was calculated. Figure 2 shows the ROC curve for the test result of this dataset.

**Figure 1.**
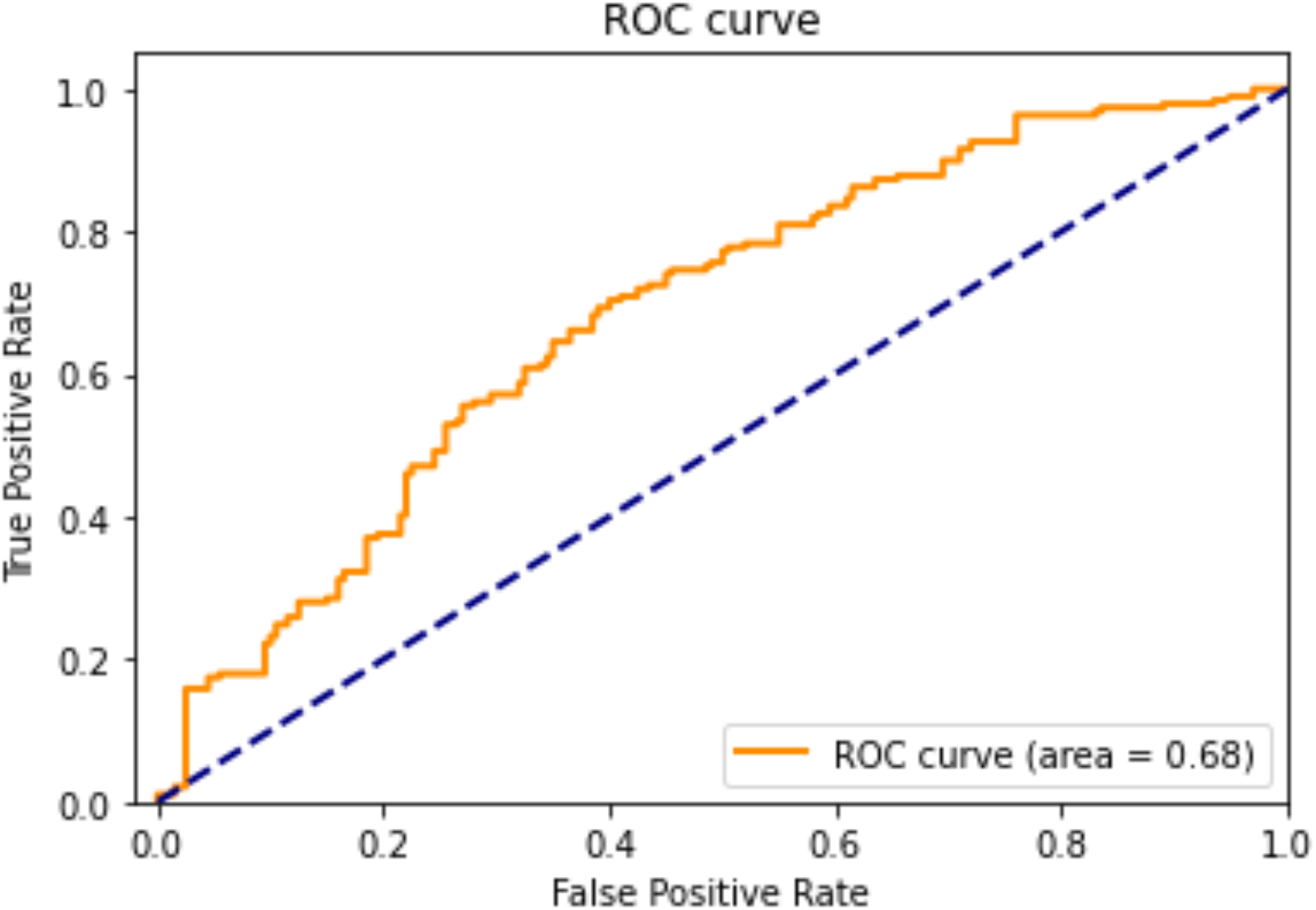
The ROC curve of prediction result to experimentally validated neoantigen peptides and negative peptides.

**Figure 2.**
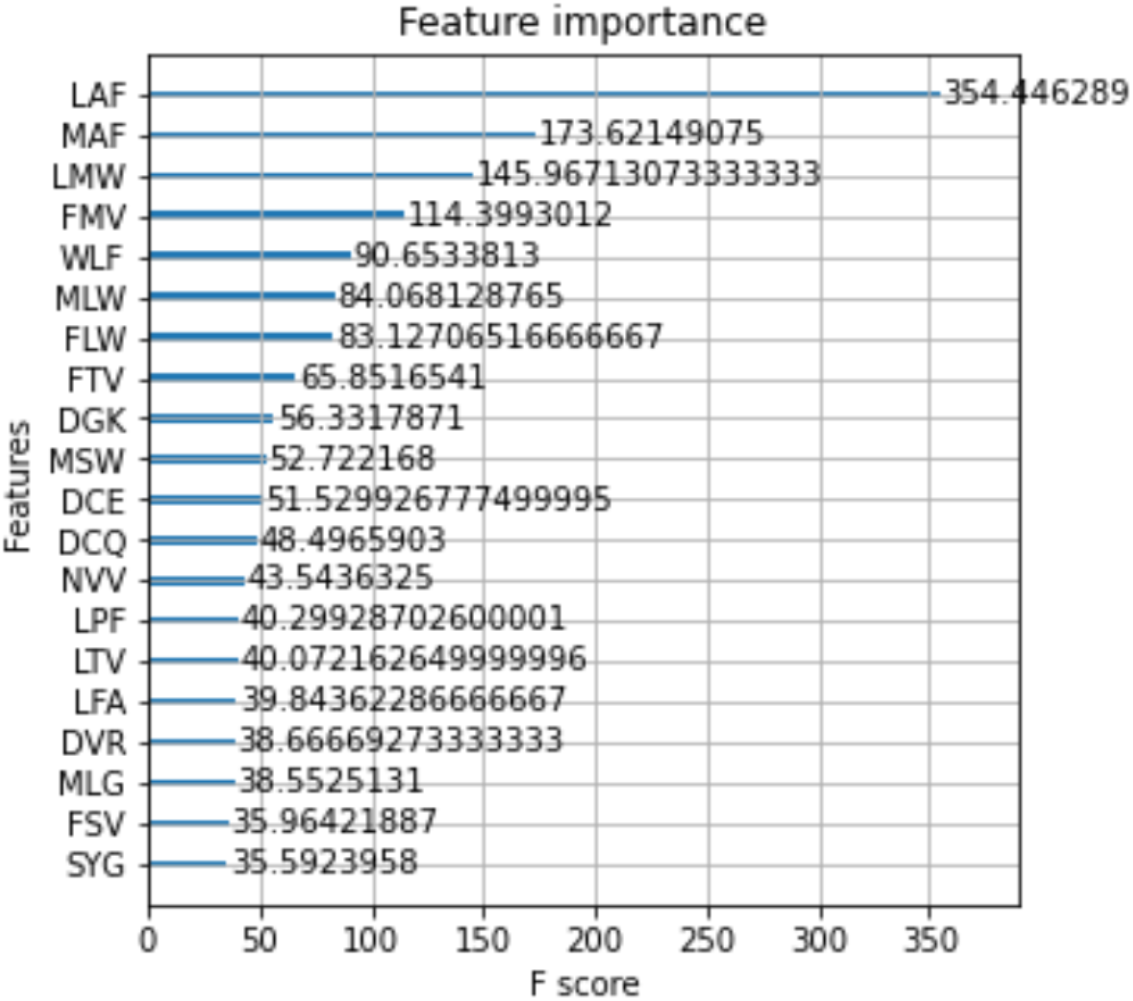
Feature importance for 20 top-ranked tri-peptides in the trained model and their F-scores.

### Top weighted tri-peptides

Our model relies on the matrix of tri-peptide from all combinations of any three amino acids. In Figure 2, we listed 20 top-ranked tri-peptides in the optimal model. The amino acids, Leucine (L), Phenylalanine (F), and tryptophan (W) frequently occur in the top 5 tri-peptides. This indicates that these three amino acids are enriched in the top-ranked tri-peptides. Therefore, these three amino acids and their combinations may play an important role in NHC binding. However, because we used the tri-peptide similarities as features and the values of these three amino acids in the substitution matrix are high, it could cause the model to favor these three amino acids.

### Examples cancer neoantigens

For example, the cancer mutant peptide, LLLMSTLGI, is in the protein coded by the gene *KLRD1* in a pancreatic cancer patient, obtained from dbPepNeo database (Tan et al., 2020). It was validated as an epitope of HLA-A*02:01 by T-cell response. Our model assigned a score of 0.753 to this mutant peptide, while giving 0.549 to the corresponding wildtype peptide, LSLMSTLGI, only one amino acid mutation. As we described before, Leucine (L) is important for our model, and therefore, for this peptide, the substitution of Serine (S) to Leucine (L) causes the peptide from a non-MHC-epitope to an MHC-epitope.

## Conclusion

The accuracy for cancer neoantigen identification based on MHC epitope prediction was increased by concurrently using tri-peptide similarity model that was trained by filtered known MHC-I epitopes. For experimentally validated cancer neoantigens, our model showed success rates from 52.2% to 72.2%, and the corresponding AUC value is 0.685. The program and all datasets are available at https://github.com/augustthedoodle/mhc_binding

## Materials and Methods

### Datasets

For the training and testing dataset, we used the datasets from NetMNCpan-4.0 (Jurtz et al., 2017), which were downloaded from http://www.cbs.dtu.dk/suppl/immunology/NetMHCpan-4.0/. We kept human MCH-I class binding peptides only (i.e. HLA-A, HLA-B, and HLA-C). A peptide is considered as a binding peptide if the smallest log50k transformed binding affinity > 0.6 whereas a peptide is considered as a negative if the smallest log50k transformed binding affinity <0.1. We kept MCH-I class binding epitopes that are 9 amino acids long. There are 3719 positive peptides and 11632 negative peptides, respectively.

We also collected four cancer neoantigen datasets for an independent test. Two datasets from dbPepNeo database (Tan et al., 2020). They are HC-neoantigens and MC-neoantigens for neoantigen information identified by T-cell response and by mass spectrometry and whole-exome sequencing, respectively. The third dataset was obtained from NEPdb (Xia et al., 2021). The fourth data is the IEDB validated cancer neoantigens downloaded from TSNAdb (Wu et al., 2018). A negative set of peptides with mutations from cancer genomes was obtained from NEPdb (Xia et al., 2021). For these datasets, only 9AA peptides were kept, and their corresponding wildtype peptides were saved as well for a comparison.

### Encoding

We used tri-peptide subsequence space to encode the features for classification. The feature vector is a space of 20^3^ attributes. For one 9-amino-acid peptide, we used a sliding window with 3 digits to get 7 3-amino-acid blocks (tri-peptides). For each tripeptide, we compare with all possible three amino acid combinations (i.e. 20^3^ types of tri-peptides), and the score is the sum of three amino acid substitution scores from blosum62 matrix. Then, all 7 by 20^3^ vectors will be summed one tri-peptide by one tri-peptide, and the final 20^3^ vector was the feature vector for the mode.

### Training/Prediction procedure for Deep learning

We used XGBoost implemented in scikit-learn as the classifier. XGBoost is an implementation of gradient boosted decision trees designed for speed and performance that is dominative competitive machine learning. For the XGBoost model, to reduce the overfitting, we adjusted hyperparameters such as maximum depth and minimum child weight. The maximum depth was set to only 6 to reduce overfitting and the minimum child weight was 1. We can use 5-fold cross-validation. This means we need to group training set into 5 groups and use 4 of them as the training and the last one for testing. The optimal parameter set for training was determined by the AUCs. The contribution of each mutation to the prediction was measured by a SHAP (SHapley Additive exPlanations) value on the test dataset, which is an alternative to permutation feature importance (Lundberg and Lee, 2017).

### Evaluation methods

The statistical terms, sensitivity (Sn), precision (P), and F-measure, are defined in the following equations:

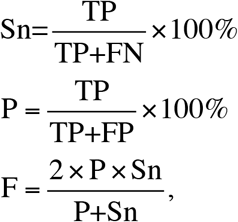

where TP, TN, FP, and FP stand for true positive, true negative, false positive, and false negative, respectively. F-measure is used to determine the optimal prediction results. The Python package, sklearn, was used to training, conduct cross-validation, and calculate the AUC.

## Acknowledgments

CZ designed the study and prepared the data. CX and YS developed the code. CZ and CX drafted the manuscript. CZ supervised the whole project. All authors read and approved the final manuscript. CX was a high-school student and conducted her summer research for this project.

## Notes

### Competing Interest Statement

The authors have declared no competing interest.

